# Gliflozins, sucrose and flavonoids are allosteric activators of lecithin:cholesterol acyltransferase

**DOI:** 10.1101/2024.06.18.599491

**Authors:** Akseli Niemelä, Laura Giorgi, Sirine Nouri, Betül Yurttaş, Khushbu Rauniyar, Michael Jeltsch, Artturi Koivuniemi

## Abstract

Lecithin:cholesterol acyltransferase (LCAT) serves as a pivotal enzyme in preserving cholesterol homeostasis via reverse cholesterol transport, a process closely associated with the onset of atherosclerosis. Impaired LCAT function can lead to severe LCAT deficiency disorders for which no pharmacological treatment exists. LCAT-based therapies, such as small molecule positive allosteric modulators (PAMs), against LCAT deficiencies and atherosclerosis hold promise, although their efficacy against atherosclerosis remains challenging. Herein we utilized a quantitative in silico metric to predict the activity of novel PAMs and tested their potencies with in vitro enzymatic assays. As predicted, sodium-glucose cotransporter 2 (SGLT2) inhibitors (gliflozins), sucrose and flavonoids activate LCAT. This has intriguing implications for the mechanism of action of gliflozins, which are commonly used in the treatment of type 2 diabetes, and for the endogenous activation of LCAT. Our results underscore the potential of molecular dynamics simulations in rational drug design.

## INTRODUCTION

Lecithin:cholesterol acyltransferase (LCAT) is a plasma enzyme which converts cholesterol into cholesteryl esters (CE) primarily while bound to high-density lipoprotein (HDL) [1]. LCAT has three domains: the membrane binding domain (MBD), which LCAT utilizes to bind to lipoproteins; the α/β-hydrolase domain, which contains the active site; and the cap domain, whose lid loop controls whether lipids can access the active site [2,3]. The lid in its open state enables a phospholipid to reach the active site and react with the SER-HIS-ASP catalytic triad, resulting in the cleavage of the fatty acid chain at the sn-2 position. If cholesterol is available the cleaved fatty acid is transferred to the free hydroxyl group of cholesterol, generating a CE molecule [1]. On nascent discoidal HDL, the formed CEs travel to the core of the disc and eventually morph it into mature spherical HDL. Thus, LCAT is essential in HDL metabolism and the associated process of cholesterol homeostasis called reverse cholesterol transport (RCT), in which the cholesterol from extrahepatic tissues is transported via HDL to the liver for elimination.

If LCAT function is impaired cholesterol accumulates into tissues causing fish-eye-disease and the more severe familial LCAT deficiency, which leads to kidney damage. Enzyme replacement therapy has shown promise against these conditions [4,5], but wider interest lies in whether LCAT activity and the underlying RCT mechanisms can be harnessed against atherosclerosis, which is caused by the build-up of arterial plaques comprised of cholesterol, fatty material, calcium and other components. Early success in in vivo trials with recombinant human LCAT has not yet translated to clinical benefit [6,7], and its atheroprotective potential has remained elusive. A factor of this is that LCAT is also active on low-density lipoproteins (LDL), which, opposite to HDL, transport cholesterol to extrahepatic tissues [8], and that simply increasing the cholesterol content of HDL may lead to passive transport of free cholesterol back to the tissues that donated it [9]. The poor efficacy of HDL cholesterol content increasing CETP inhibitor anacetrapib against major cardiovascular events supports this [10], and focus has shifted from HDL cholesterol to cholesterol efflux capacity [11]. However, even reconstituted HDL particles, whose mechanism of action is to act as a cholesterol acceptor, were not effective enough in clinical trials to warrant commercialization [12]. It is important to note that the CETP inhibitor anacetrapib attained more pronounced results during an extended follow-up period [13], so RCT based therapies may require long-term use for clinical benefit. For conducting long-term clinical trials, orally deliverable small molecules are most suitable due to their ease of administration and cost-effectiveness.

DS-8190a is a small molecule positive allosteric modulator (PAM) of LCAT which showed promise in vivo [14]. Manthei et al. investigated a set of its predecessors and showed that they allosterically boost the activity of LCAT by binding to the MBD [15]. Molecular dynamics (MD) simulations of this set with LCAT revealed a conformational change they induce, where the MBD spatially shifts away from the lid and its binding cavity [16]. This repositioning facilitates the proper opening of the lid and active site, potentially improving the access of lipids into the active site and or their orientation for catalytic activity.

Herein we describe an in silico model for the existing positive allosteric modulators of LCAT that predicts their capability to activate LCAT quantitatively. We conducted an in silico screen with FDA approved compounds and plant secondary metabolites and subsequently employed the model with the aim of finding novel LCAT activating compounds and validating our quantitative metric with a different set of small molecules. As improved LCAT activity would likely have innocuous or benign effects on cholesterol distribution, we hypothesized that known FDA approved compounds may possess unknown LCAT activating properties. Secondary metabolites were included in the screen as we had noted the scaffold similarity between DS-8190a and flavonoids.

The in silico screen predicted that gliflozins, sucrose and flavonoids would bind and activate LCAT, while mebendazole would bind but not activate the enzyme. Phospholipase and acyltransferase assays with LCAT confirmed that our predicted allosteric activators promote the activity of LCAT in vitro within the predicted levels. Our findings have interesting implications for how reverse cholesterol transport is linked with diet and the mechanism of action of gliflozins, which are one of the primary pharmacological tools for the treatment of type 2 diabetes.

## RESULTS

### In silico screens

We had previously performed molecular dynamics (MD) simulations to a set of Daiichi Sankyo based allosteric activators of LCAT [16]. By analysing these trajectories again we discovered that the CYS50-ASN65 α-carbon distance correlated strongly with the activity the compounds induce in LCAT in a 4-methylumbelliferyl palmitate (MUP) based phospholipase assay (Fig. 1A-C) [15]. We set out to perform a combined docking and MD screen to find activators in plant secondary metabolites and FDA approved compounds (n=175 and n=1615 respectively) and to predict their activity with the CYS50-ASN65 distance (Supplementary Table 1).

**Fig. 1.**
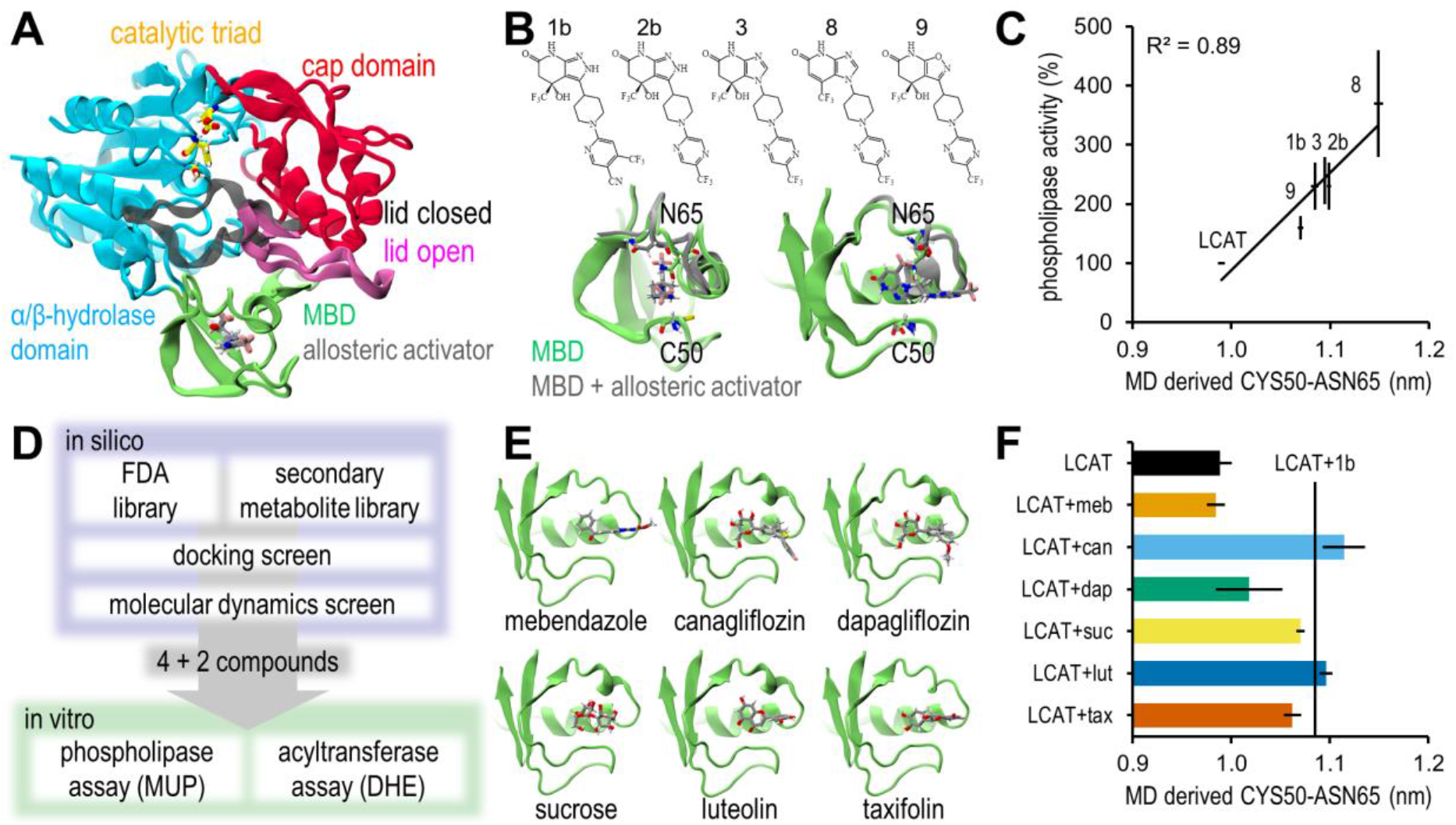
In silico results. (A) Domains of LCAT with its lid in open and closed states based on PDB ID 6MVD and 5TXF respectively [3,15]. (B) Chemical structures of the allosteric activators of Daiichi Sankyo and their effect on the MBD. Specifically, ASN65 shifts away from CYS50. (C) Correlation between the in vitro phospholipase activities and molecular dynamics derived CYS50-ASN65 α-carbon distance. In vitro mean and SEM results were collected from ref [15]. MD data reported as mean and SEM of 1 µs simulations split to ten 100 ns sequences. (D) Overview of the in silico screen. (E) Compounds discovered from the in silico screen and their binding poses in the MBD. (F) The discovered compounds’ induced CYS50-ASN65 distances. The mean distance of LCAT with 1b is marked (230 % activity compared to LCAT). Data reported as mean and SEM of five replicate 200 ns simulations (last 100 ns analysed).

Docking simulations with the Daiichi Sankyo based activators revealed docking scores in the range of -5.2 - -7.2 (Supplementary Table 2). Thus a score of -7.0 was deemed a suitable cut-off for LCAT binding. Out of the docking screen 10 FDA approved compounds and 5 secondary metabolites were screened with molecular dynamics simulations (Supplementary Table 3). The best binding secondary metabolites were flavonoids. Compounds which did not stay in the docked pose were rejected in preliminary molecular dynamics simulations, and out of the remaining compounds canagliflozin, dapagliflozin, sucrose, luteolin and taxifolin were selected as activators of LCAT, and mebendazole as a negative control for the in vitro assays (Fig. 1D-F). Five replicate simulations of 200 ns were performed for this set, whose last 100 ns were analysed to gain an accurate prediction for activity. Canagliflozin, dapagliflozin and sucrose used their glucose units to bind to the MBD, and presented with a tendency to reorient during the replicate simulations (one, three and one out of five simulations respectively) (Supplementary Fig. 1). Despite this, the reorientation did not consistently result in different CYS50-ASN65 distances, although dapagliflozin presented with markedly higher variability (Supplementary Table 2).

### In vitro assays

Based on the in silico results canagliflozin, dapagliflozin, sucrose, luteolin and taxifolin were selected for in vitro validation as possible new activators of LCAT, while mebendazole was chosen as negative control. The phospholipase activity of LCAT in the presence of the selected compounds was determined with a MUP based phospholipase assay (Table 1, Fig. 2A). Supplementary Table 4 shows Tukey’s test groupings of each concentration point. As predicted, mebendazole did not enhance the catalytic activity of LCAT, showing an identical reaction rate curve compared to LCAT alone, whereas canagliflozin, dapagliflozin, sucrose, luteolin and taxifolin improved the reaction rate at different MUP concentrations. Particularly canagliflozin, dapagliflozin and taxifolin increased V_max_ by 140 %, 80 % and 70 % respectively, and canagliflozin and sucrose decreased K_m_ by 60 % and 80 % respectively. Phospholipase efficiency, which takes into consideration V_max_, K_m_ and LCAT concentration, was improved 540 % by canagliflozin.

**Table 1.** In vitro enzyme kinetics results. Phospholipase activity was measured with a MUP phospholipase assay and acyltransferase activity with a DHE esterification assay. Data reported as mean±SEM.

| configuration | phospholipase $V_{\max}$ ($\mu\text{M h}^{-1}$) | phospholipase $K_m$ ($\mu\text{M}$) | acyltransferase $V_{\max}$ ($\mu\text{M h}^{-1}$) | acyltransferase $K_m$ ($\mu\text{M}$) |
| --- | --- | --- | --- | --- |
| LCAT | 6.33±0.47 | 52.82±7.96 | 8.60±0.28 | 11.25±0.71 |
| LCAT+mebendazole | 7.00±0.24 | 57.39±3.32 | 9.04±2.09 | 23.25±10.45 |
| LCAT+canagliflozin | 15.27±1.20 | 20.01±4.70 | 14.73±0.81 | 8.01±1.47 |
| LCAT+dapagliflozin | 11.48±0.93 | 43.20±11.40 | 12.38±0.71 | 9.50±2.03 |
| LCAT+sucrose | 7.41±1.43 | 11.86±6.99 | 11.49±0.36 | 8.61±0.67 |
| LCAT+luteolin | 9.35±1.57 | 24.18±11.29 | 11.52±0.63 | 7.92±1.07 |
| LCAT+taxifolin | 10.64±0.09 | 26.49±0.57 | 14.44±1.50 | 15.64±2.95 |

**Fig. 2.**
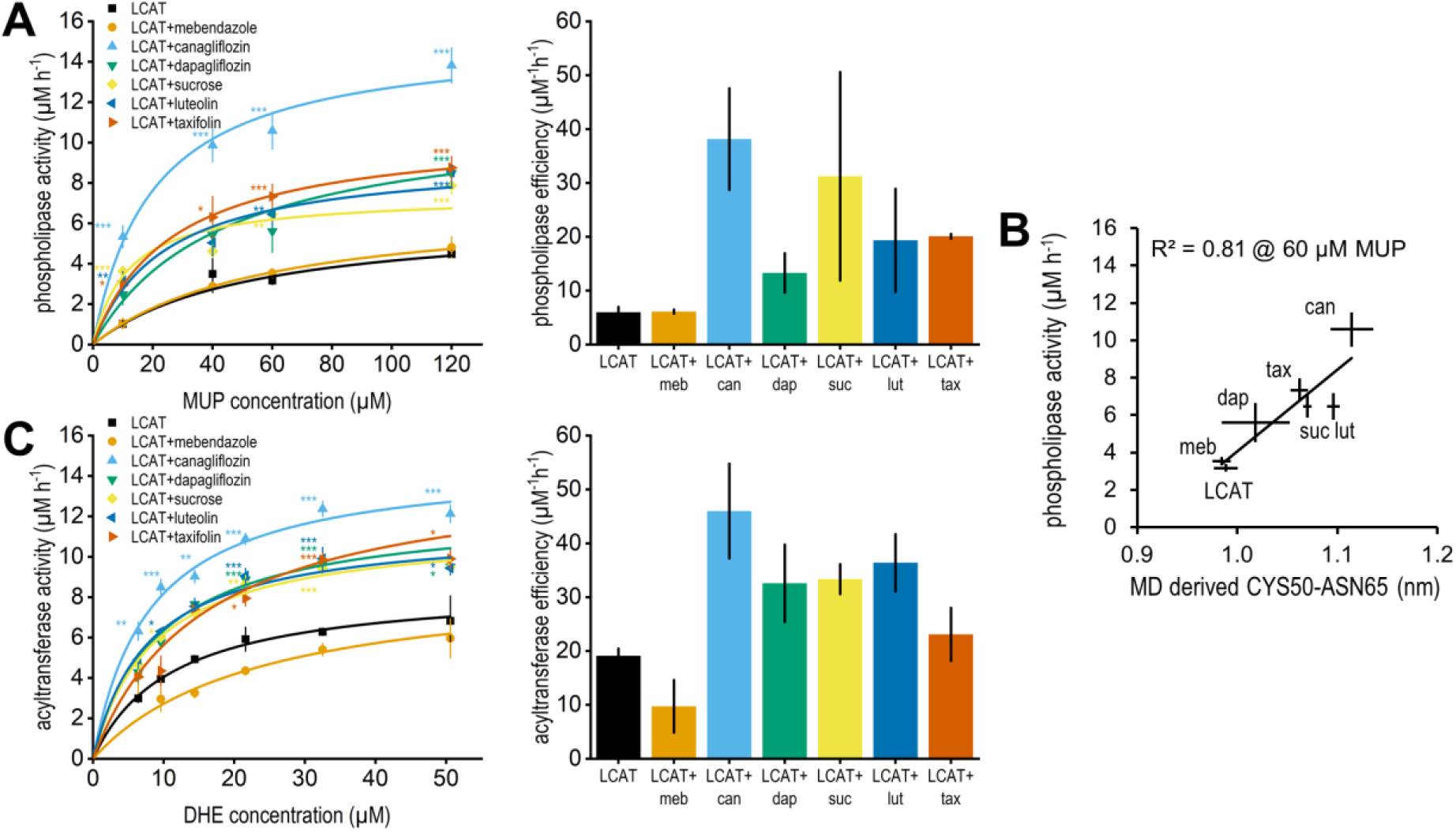
In vitro activity results. (A) Phospholipase activity rate (left) and catalytic efficiency (right). Data reported as mean and SEM of triplicate measurements. Catalytic efficiency was calculated by dividing k_cat_ (V_max_ divided by enzyme concentration) with K_m_. Error bars are SEM attained by propagating the SEMs of V_max_ and K_m_. Statistical significance between all configurations at all concentration points were calculated with one-way ANOVA followed by Tukey’s test. Differences to LCAT are shown with the following levels: *p<0.05, **p<0.01, ***p<0.001. (B) Correlation between the in vitro phospholipase activities at 60 µM MUP concentration plotted with the in silico CYS50-ASN65 distances. MD data reported as mean and SEM of five replicate 200 ns simulations (last 100 ns analysed). (C) Acyltransferase activity rate (left) and catalytic efficiency (right).

Our in silico predictive model was based on a MUP phospholipase assay where the LCAT:MUP ratio was ∼1:2700, and the LCAT:PAM ratio was increased until ∼1:1600 [15]. Thus, we used the phospholipase activity values at 60 µM MUP (1:200:3000 LCAT:PAM:MUP ratio) to validate our predictive model (Fig. 2B). Since our MUP assay was different, a new slope was fitted and the correlation remained high (R^2^=0.81). A decrease in the correlation was not unexpected given that the true binding poses are unknown, more variability in the poses was present and our LCAT:PAM ratio was smaller. Pearson correlation coefficients between all in vitro measurements and the CYS50-ASN65 distances are shown in Supplementary Table 5.

The acyltransferase activity of LCAT was determined with a synthetic HDL (sHDL) based assay, where fluorescent cholesterol-like dehydroergosterol (DHE) was incorporated into the sHDL. The results were in line with the phospholipase assay (Table 1, Fig. 2C). In terms of V_max_, particularly canagliflozin and taxifolin increased it by 70 %. Canagliflozin maintained an acyltransferase efficiency improvement of 140 %.

## DISCUSSION

Through molecular dynamics (MD) simulations we had previously identified that allosteric activators of LCAT increase the MET66-ILE231 α-carbon distance [16]. A reanalysis of these simulations revealed that their LCAT promoting activities correlated strongly with their capability to increase the CYS50-ASN65 α-carbon distance. As the distance increases, the MBD of LCAT shifts away from the lid cavity, which possibly enhances the lid’s capability to change states, alleviates lipid entry to the active site, and or stabilises the open state of the lid. We set out to discover allosteric activators of LCAT from known FDA approved compounds and secondary metabolites with a virtual screen and utilized the distance metric to predict their activities.

In vitro assays confirmed that sodium-glucose cotransporter 2 (SGLT2) inhibitors canagliflozin and dapagliflozin, sucrose, and flavonoids luteolin and taxifolin increased LCAT’s activity to predicted levels. Our negative control, the anthelmintic mebendazole, could not activate LCAT. As far as we are aware, this is the first time molecular dynamics has revealed a compound induced conformational change which is quantitatively linked with activity. Although passable binding poses for novel compounds can be attained via docking, we believe this method would be most useful during lead optimization with validated binding poses, since in the timescales attainable by MD ligands can drive the receptor conformation into thermodynamically stable local minimas irrelevant in the timescales of activity assays. In other words, MD derived predictions are useful only if one is sampling the correct conformation of the ligand-receptor complex. A limitation of our study is that the binding poses are not experimentally validated, which contributed to the variability of dapagliflozin’s CYS50-ASN65 distances. Similarly whether mebendazole is able to bind to the MBD in vitro is unknown, but the inhibitive effect in the acyltransferase assay in all but one DHE concentration suggests allosteric modulation occurs.

Sucrose and the gliflozins utilized a glucose unit to bind to the MBD of LCAT. This suggests that the MBD may be a more general sugar binding domain that LCAT could use to interface with glycans. Apolipoprotein A1, the most potent cofactor of LCAT on lipoprotein particles, is predicted to have two glycosylation sites [17], and intriguingly, LCAT has poorer activity on desialylated HDL [18]. However, this has been attributed to the reduced electronegative charge [19]. Speculatively, lipoprotein glycans could either improve LCAT activity by facilitating lid opening, or inhibit it by preventing access to the lipid surface. Further studies are required to elucidate whether the MBD can accommodate a glycan chain in solution or at the surface of lipoprotein particles, and whether this affects LCAT’s localization on the particles or distribution in plasma. LCAT activity measured in the blood of diabetes patients before and after insulin treatment confirmed that glucose on its own cannot activate LCAT [20].

Gliflozins are FDA approved compounds for the treatment of type 2 diabetes. They decrease plasma glucose levels by inhibiting SGLT2, which would otherwise reabsorb almost all glucose from renal filtrate. Based on in vivo experiments with allosteric activators and clinical trials with recombinant human LCAT, an increase in LCAT activity is expected to increase both HDL and LDL cholesterol [14,21,22]. A meta-analysis of 48 randomized controlled trials confirms that canagliflozin and dapagliflozin elicit these changes, with canagliflozin having a stronger effect on LDL cholesterol [23]. Although this is consistent with our LCAT activity assay results, SGLT2 inhibitors have been proposed to affect lipoprotein cholesterol levels through different mechanisms, such as hemoconcentration and downregulation of LDL receptors [24]. It is also important to note that 99 % of canagliflozin and 91 % of dapagliflozin is bound to proteins in plasma, which reduces the fraction available to bind to LCAT.

In different mouse and rat models luteolin has been reported to either increase or not affect HDL cholesterol and to either decrease or not affect LDL cholesterol [25,26,27,28]. Taxifolin has been reported to either increase HDL cholesterol or decrease it with a cholesterol free diet and to decrease LDL cholesterol [29,30,31]. However, flavonoids affect many pathways in the body, and accordingly these studies present a variety of LCAT independent mechanisms that explain the changes in lipoprotein cholesterol levels [32], so it cannot be determined whether LCAT activation contributed. Furthermore, like sucrose, flavonoids have poor oral bioavailability which reduces the potential clinical benefit they may bring via LCAT activation [33,34].

There are other LCAT-like enzymes whose MBDs might accept allosteric modulators. The MBD of human lysosomal phospholipase A2 shares a sequence identity of 56 % with the MBD of LCAT (residues 36-99) and a similar fold (UniProt ID Q8NCC3) [2]. LCAT-like phospholipase A in *Arabidopsis thaliana* has a sequence identity of 32 %, although its AlphaFold predicted MBD fold is more deformed (UniProt ID Q9FZI8) [35,36]. Phospholipases have a wide variety of functions in metabolism and signal transduction [37]. Speculatively, perhaps LCAT’s capability to be activated by flavonoids originates from its progenitors in plants. The recent advances in clustering AlphaFold predicted protein structures across species would enable a deeper study into the evolution of allosteric modulation of LCAT-like enzymes [38]. Human LCAT (UniProt ID P04180) belongs to the cluster whose lowest common ancestor is an uncharacterized protein of Polyangiaceae bacterium (UniProt ID A0A850BM54), which lacks the binding site defining bottom loop of the MBD.

In conclusion, we present a conformational change allosteric activators of LCAT induce in molecular dynamics simulations that can be used to predict their in vitro activities. With this model we discovered that antidiabetic SGLT2 inhibitors canagliflozin and dapagliflozin possess a clinically novel potentially atherosclerosis preventing mechanism of action: the allosteric activation of LCAT. We also discovered that flavonoids luteolin and taxifolin possess this same mechanism of action. Further studies are required to show whether a wider variety of flavonoids share this trait, and whether the consumption of dietary flavonoids results in clinically relevant improvements in the lipid profile via LCAT activation.

## METHODS

### Discovery of activity predicting model

We reanalysed our previously simulated trajectories of compounds 1b, 2a, 2b, 3, 6, 8 and 9 [16]. Compounds 2a and 6 were left out of the analysis since the hybridization state of 2a was not accommodated by the MBD properly, resulting in the bottom loop unfolding, and since 6 could not bind to the MBD in vitro [15]. We discovered that the activity the compounds induce in vitro correlated with their capability to shift the top loop of the MBD relative to the bottom loop in silico. The α-carbon distance between CYS50 and ASN65 was found to be the best marker.

### Collection of virtual compound libraries

Two virtual compound libraries were collected. The first was the complete FDA approved drug library from the ZINC15-database (n=1615) [39]. The second of plant secondary metabolites was attained by a systematic literature review (n=175) in June 2021. The Scopus database was filtered first with keywords LCAT or “lecithin cholesterol acyltransferase” and with the name of a secondary metabolite. Search was performed within article titles, abstracts and keywords. LCAT was added to the filter to find compounds which affect LCAT expression or cholesterol levels. The names for secondary metabolites were attained from ref [40]. If this produced over 50 articles, an additional filter was added with keywords for natural products. Finally, articles were filtered by titles and abstracts. The resulting primary publications were crudely scanned for compounds. If the articles only had a compound source listed, such as a specific plant, secondary articles were searched for the secondary metabolites of that plant. The coordinate files for these compounds were attained from PubChem [41]. The articles and their filtering are shown in detail in Supplementary Tables 6-8.

### Docking screen

The docking screen was performed with Schrödinger software (Release 2020-3. Schrödinger, LLC, New York, 2021). All compounds were first subjected to LigPrep with standard settings, except ionization states were generated for pH 7.4 ± 2.0 and no tautomers were generated. If the coordinate file of the compound had uncertain stereochemistry, LigPrep generates all possible states. Next, as out of the compounds tested in ref [15] the charged compound 6 was the only one unable to bind to LCAT in vitro, all compounds with any charged groups were removed with LigFilter. Compounds with a molecular mass below 100 g/mol or above 600 g/mol were also removed. The MBD of LCAT, using the crystal structure solved with compound 1b bound to the MBD (PDB ID 6MVD) [15], was prepared for docking by picking compound 1b as the model ligand and using standard settings. Finally, docking with standard precision (SP) using Glide [42] was performed on both libraries, and the resulting compounds with docking scores below -7 were docked again with extra precision (XP). Compounds 1b, 2b, 3, 8 and 9 were also XP-docked for comparison. Standard settings were used, except only the most likely conformation of each compound was retained. The FDA approved compounds were ranked according to docking score and manually picked to ensure a wide range of scaffolds. All of the best binding compounds from the secondary metabolite library were flavonoids, so a representative sample of compounds with different binding poses was picked.

### Molecular dynamics screen

The same system configuration was used as in our previous study [16]. To parametrize the compounds the Amber18 modeling package’s antechamber software was utilized [43]. The compounds were geometry optimized and the partial charges were determined with the Gaussian software (16 revision A.03) utilizing the Hartree-Fock method with the 6-31G* basis set. The Merz-Kollman scheme was used to determine the electrostatic potentials around the molecules which were then derived to partial charges with the restrained electrostatic potential method. The GAFF force field was used to describe the Lennard-Jones and bonded parameters [44]. For LCAT, the AMBER99SB-ILDNS force field was used [45], and for water the TIP3P model.

The GROMACS simulation package (Version 2020.2) was used to run all MD simulation systems [46]. LCAT was placed to the centre of an 8 x 8 x 8 nm^3^ box with ∼14600 water molecules. The compounds were placed into the MBD of LCAT in the binding pose discovered with docking, and before production runs the systems were energy minimized with the steepest descent algorithm. With the v-rescale thermostat, a 310 K temperature was used with a 0.5 ps coupling constant [47]. And with the Parrinello-Rahman barostat, a 1.013 bar isotropic pressure was used with a 10 ps coupling constant. A 1.0 nm cut-off was used for the Particle-Mesh Ewald summation scheme and Lennard-Jones interactions [48].

The systems were first run for 1 µs. Any compound that dissociated or whose binding pose changed markedly was rejected. Mebendazole, canagliflozin, dapagliflozin, sucrose, luteolin and taxifolin were chosen for in vitro assays. Upon running duplicates it was noted that the compounds binding with a glucose unit, sucrose, canagliflozin and dapagliflozin, could shift their binding pose in the MBD. Thus, we decided to run an additional equilibration for this set with all heavy LCAT atoms restrained with 1000 kJ/mol nm^2^ in x-, y- and z-direction for 100 ps and to run this set without restraints for 200 ns with five replicates. Same equilibration and production runs were performed for LCAT without compounds. The last 100 ns of the trajectories were analysed with the GROMACS tool gmx mindist to calculate the mean CYS50-ASN65 α-carbon distance.

### Materials for experiments

Sucrose, mebendazole, taxifolin, 4-methylumbelliferone (MU), and cholesterol oxidase (COx) were obtained from Sigma-Aldrich (Merck, Darmstadt, Germany). Canagliflozin, dapagliflozin, luteolin and 4-methylumbelliferyl palmitate (MUP) were purchased from Cayman Chemical Company (MI, USA). 22A peptide (PVLDLFRELLNELLEALKQKLK) was obtained from Peptide Protein Research Ltd. (Fareham, UK). 1,2-Dimyristoyl-sn-glycero-3-phosphocholine (DMPC) and dehydroergosterol (DHE) were purchased from Avanti Polar Lipids Inc. (AL, USA).

### Cell culture, protein production and purification

Recombinant LCAT was produced in 293T cells (ATCC CRL-3216). A synthetic human codon-optimized cDNA coding for LCAT with a C-terminal 6x histidine-tag in pCHOKE-B vector (GenBank PP915207) was transfected into 293T cells. Cells were selected with zeocin (75 ng/mL, Invivogen) grown in semi-suspension in 150 mm plates in a mixture of serum free media (ACF CHO medium EX-CELL Merck, CHO ONE media Capricorn Scientific, OptiMEM-CHO Gibco), supplemented with 1% Primatone (Merck), 0.7% Glutamine (Gibco), 100 U/mL penicillin (ThermoFisher Scientific, 100 U/mL streptomycin (ThermoFisher Scientific). The media was harvested every 5 days. LCAT expression in 293T cells was verified by western blot with an anti-His antibody (Qiagen Penta-His Antibody). LCAT was purified by Ni Sepharose excel (Cytiva) immobilized metal affinity chromatography (IMAC) and polished via Superdex 200 (Cytiva) size exclusion liquid chromatography (NGC, Bio-Rad).

### Phospholipase assay

The phospholipase activity of LCAT was evaluated by using MUP as a substrate. The reaction was monitored with the increased fluorescence signal due to the formation of the reaction product MU. The assay was performed at room temperature in reaction buffer (0.1M sodium phosphate buffer, pH 7.4). 55 μL of LCAT in reaction buffer (0.01 mg/mL) were incubated at 37°C gently shaken with 0.41 μL of PAM (2 mg/mL in DMSO) or DMSO. After 1 h incubation, 2 μL of each reaction (LCAT + DMSO, LCAT + PAM, reaction buffer + PAM) was added to a black 384 well microplate, low volume (Corning). A MUP stock solution 1 mg/mL in DMSO + 3% Triton X-100 was diluted in a reaction buffer to 0, 10, 40, 60 and 120 μM. 18 μL of each dilution was added to the plate to start the reaction. The MUP stock solution was diluted immediately before use due to the self-hydrolysis of MUP to MU in aqueous buffers. The phospholipase activity of LCAT was monitored by fluorescence signal using Varioskan LUX (Thermo Scientific) plate reader at an excitation wavelength of 340 nm and an emission wavelength of 460 nm every 10 min for 60 min. Background signal due to MUP self-hydrolysis was subtracted from each result. Reactions were performed in triplicates with three independent experiments. Reaction rates were obtained by dividing the data by the assay time. To calculate the amount of MU produced in each reaction, a standard curve was generated from a MU stock solution 100 mM in DMSO and diluted further in reaction buffer. The phospholipase activity (MU mM) was plotted as a function of the MUP concentration. Data were fitted to the Michaelis-Menten kinetic equation by non-linear regression. Apparent V_max_ and K_m_ were calculated with OriginPro2024 (MA, USA) software, while phospholipase efficiency of each PAM was calculated by dividing k_cat_ by K_m_. The absence of fluorescence interference coming from PAMs was checked at an excitation wavelength of 340 nm before starting the assay (Supplementary Fig. 2). The phospholipase activities of our PAMs were compared to a positive control compound A (Supplementary Fig. 3) [49].

### Acyltransferase assay

The acyltransferase activity of LCAT was measured by using the fluorescent sterol DHE as a substrate. DHE is integrated in the structure of discoidal synthetic HDL (sHDL) made of peptides (22A) and phospholipids (DMPC). As a consequence of the acyltransferase activity of LCAT, DHE is esterified and stored in the inner core of sHDL as DHE-ester. The esterification is responsible for a shift in the emission wavelength of DHE from 372 nm to 425 nm, allowing the exclusive detection of DHE-ester and monitoring of the reaction.

DHE-sHDL was prepared via thermal cycling. In a glass vial, DHE and DMPC dissolved in chloroform were mixed with a molar ratio of 1:9. The organic solvent was dried with Rotavapor R-200 (BUCHI, Switzerland) at room temperature. The resulting dried thin lipid layer was hydrated with assay buffer (PBS 20 mM + EDTA 1 mM) and vortexed until complete dissolution. The solution was homogenized in a water bath sonicator at room temperature until transparency. 22A peptides dissolved in assay buffer were added to the vial (lipid to protein 2:1 w/w). The solution was incubated via three heating-cooling cycles (50 °C x 3 min; 4 °C x 3 min). The final concentration of DHE in the sample was 125 μM. The acyltransferase activity assay was performed in 384 well low volume black microplates with a final volume of 25 μL. PAM (0.1 mg/mL in DMSO) or DMSO were incubated with LCAT (5 μg/mL in assay buffer + 60 μM albumin (Merck)) 1:100 v/v at 37°C gently shaken. After 30 min incubation, 10 μL of each reaction (LCAT + DMSO or LCAT + PAM) was added to the plate. To start the reaction, 10 μL of DHE-sHDL solution were added in the wells. The plate was incubated for 1 h at 37 °C gently shaken. Reactions were stopped by the addition of 5 μL of stop solution (assay buffer + 5 units/mL cholesterol oxidase COx + 7% Triton X-100). Plates went through a second round of incubation at 37 °C gently shaken for 1 h to extinguish the residual fluorescence of unesterified DHE. Following the addition of LCAT + PAM/DMSO and stop solution, DHE concentration of the different sample dilutions in the wells resulted 0, 6.4, 9.6, 14.4, 21.6, 32.5 and 50.6 μM; while the final LCAT and PAMs concentration resulted respectively 2 μg/mL and 0.4 μg/mL. DHE-ester fluorescence was detected at an excitation wavelength of 325 nm and an emission wavelength of 425 nm using Varioskan LUX. Reactions without LCAT were used for background subtraction, while reactions without LCAT and stop solution without COx were used to generate a standard curve for DHE (λex 325 nm, λem 372 nm). Reactions were performed in triplicates with three independent experiments. Due to the light sensitivity of DHE, samples were protected from light during the entire preparation and assay. Data was processed via background subtraction, divided by the slope of the standard curve to obtain the amount of DHE-esters in every well and then divided further by the reaction time to determine the rate. The acyltransferase activity (DHE-ester formed) was plotted as a function of the DHE concentration. Data were fitted to the Michaelis-Menten kinetic equation by non-linear regression. Apparent V_max_ and K_m_ were calculated using OriginPro2024 software, while acyltransferase efficiency of each PAM was calculated by dividing k_cat_ by K_m_. The absence of fluorescence interference coming from PAMs was checked at an excitation wavelength of 325 nm before starting the assay (Supplementary Fig. 2). The acyltransferase activities of our PAMs were compared to the reference compound A (Supplementary Fig. 3).

### Statistical analysis

Statistical analysis for triplicate measurements was performed with a one-way analysis of variance (ANOVA) followed by Tukey’s test, using OriginPro2024 software. All data are presented as mean and SEM. A significance level of p<0.05 was used to group the configurations at the same concentration points.

## AUTHOR CONTRIBUTIONS

A.N. and L.G. contributed equally to this work. A.K., A.N. and L.G. designed the study. A.N. performed in silico screening and simulations. L.G. performed the LCAT activity experiments. L.G., A.N. and S.N. analysed and interpreted the experimental data. L.G., B.Y., K.R., and M.J. expressed and purified the LCAT enzyme. A.N. and L.G. wrote the paper with input from all authors.

## ACKNOWLEDGMENTS

A.K. discloses support for the research of this work from the Scientific Council for Biosciences, Health and the Environment of the Research Council of Finland [350636]. The authors wish to acknowledge CSC–IT Center for Science, Finland, for computational resources.

