## Supplementary Information for "Gliflozins, sucrose and flavonoids are allosteric activators of lecithin:cholesterol acyltransferase"

**Supplementary Table 1.** The number of compounds at the end of each step of the in silico screen.

**Supplementary Table 2.** Docking scores and MD derived CYS50-ASN65 distances. MD data of Daiichi Sankyo compounds reported as mean and SEM of 1 µs simulations split to ten 100 ns sequences.

**Supplementary Table 3.** Preliminary MD screen results. Result meanings: + activates; - does not activate; NA not applicable, binding pose changed remarkably or dissociated.

**Supplementary Table 4.** Statistical analysis of phospholipase and acyltransferase activity results. The activity rates at each concentration point (n=3) were compared with one-way ANOVA with a post-hoc Tukey’s test. Configurations were grouped at a significance level of p<0.05. Those that do not share a letter are significantly different, and those that are significantly different from LCAT are bolded.

**Supplementary Table 5.** Correlation between MD derived CYS50-ASN65 distances and in vitro attained values.

**Supplementary Table 6.** The number of articles that passed each filter for secondary metabolite compound library. Filter 2 was used if over 50 articles passed filter 1.

**Supplementary Table 7.** Primary articles for secondary metabolite compound library without duplicates.

**Supplementary Table 8.** Secondary articles for secondary metabolite compound library.

**Supplementary Fig. 1.** Start and end conformations of the MBD and compounds of the 5 replicate MD simulations. Starting conformation is the docked pose after 100 ps MD equilibration with LCAT heavy atoms restrained. Sim 3 of sucrose, sim 1 of canagliflozin and sims 1-3 of dapagliflozin have a sideways binding pose (parallel to perspective).

**Supplementary Fig. 2.** PAMs show no background interference. Fluorescence emission spectra of our PAMs and MUP in assay buffer following excitation at 340 nm (left). MUP in aqueous buffer has a clear peak at 460 nm. Fluorescence emission spectra of our PAMs and DHE in assay buffer following excitation at 325 nm (right). DHE in aqueous buffer has a peak at 372 nm, while DHE-ester has a peak at 425 nm.

**Supplementary Fig. 3.** Assay validation with positive control compound A. Phospholipase activity at 60 µM MUP (left). Acyltransferase activity at 32.5 µM DHE (right).

**Supplementary Table 1**

| in silico screen step | secondary metabolite library | FDA approved library |
| --- | --- | --- |
| complete library | 175 | 1615 |
| LigPrep | 623 | 2586 |
| Ligfilter | 200 | 883 |
| Glide SP docking | 198 | 872 |
| docking score < -7 | 32 | 41 |
| Glide XP docking | 32 | 41 |
| preliminary MD | 5 | 10 |
| 5 replicates MD | 2 | 4 |
| in vitro screen | 2 | 4 |

**Supplementary Table 2**

|  |  | MD CYS50-ASN65 distance | | | | |
| --- | --- | --- | --- | --- | --- | --- |
| compound | docking score | sim 1 | sim 2 | sim 3 | sim 4 | sim 5 |
| - | - | 0.99±0.004 | - | - | - | - |
| 1b | -7.2 | 1.08±0.004 | - | - | - | - |
| 2b | -6.7 | 1.10±0.002 | - | - | - | - |
| 3 | -6.2 | 1.09±0.002 | - | - | - | - |
| 8 | -5.2 | 1.15±0.004 | - | - | - | - |
| 9 | -6.1 | 1.07±0.003 | - | - | - | - |
| - | - | 0.97 | 1.02 | 1.00 | 0.94 | 1.01 |
| mebendazole | -7.1 | 0.99 | 0.97 | 0.98 | 1.02 | 0.96 |
| canagliflozin | -8.2 | 1.04 | 1.14 | 1.18 | 1.08 | 1.13 |
| dapagliflozin | -7.9 | 0.94 | 1.06 | 1.06 | 1.12 | 0.92 |
| sucrose | -10.7 | 1.07 | 1.09 | 1.06 | 1.06 | 1.06 |
| luteolin | -8.8 | 1.08 | 1.11 | 1.11 | 1.09 | 1.09 |
| taxifolin | -9.0 | 1.06 | 1.03 | 1.09 | 1.08 | 1.06 |

**Supplementary Table 3**

| compound library | ID | name | preliminary MD screen result |
| --- | --- | --- | --- |
| FDA approved | 1 | sucrose | + |
|  | 2 | kuvan (ZINC ID: ZINC2539827) | NA |
|  | 3 | inosine | NA |
|  | 4 | brilinta | NA |
|  | 5 | canagliflozin | + |
|  | 6 | dapagliflozin | + |
|  | 7 | piceid | NA |
|  | 8 | thymidine | + |
|  | 9 | kuvan (ZINC ID: ZINC4228257) | NA |
|  | 10 | mebendazole | - |
| secondary metabolite | 1 | diosmin | NA |
|  | 2 | taxifolin | + |
|  | 3 | luteolin | + |
|  | 4 | leucocyanidin | NA |
|  | 5 | catechin-7-O-glucoside (PubChem ID: 44257085) | NA |

**Supplementary Table 4**

|  |  | LCAT | LCAT+  meb | LCAT+  can | LCAT+  dap | LCAT+  suc | LCAT+  lut | LCAT+  tax |
| --- | --- | --- | --- | --- | --- | --- | --- | --- |
| phospholipase MUP concentration (µM) | 10 | C | C | **A** | C,B | **B** | **B** | **B** |
|  | 40 | C,D | D | **A** | B,C | B,C,D | B,C,D | **B** |
|  | 60 | C | C | **A** | B,C | **B** | **B** | **B** |
|  | 120 | C | C | **A** | **B** | **B** | **B** | **B** |
| acyltransferase DHE concentration (µM) | 6.4 | B | B | **A** | A,B | A,B | A,B | B |
|  | 9.6 | C,D | D | **A** | A,B,C | **A,B** | **A,B** | B,C,D |
|  | 14.4 | B,C | C | **A** | A,B | A,B | A,B | A,B |
|  | 21.6 | C | C | **A** | **B** | **B** | **B** | **B** |
|  | 32.5 | C | C | **A** | **B** | **B** | **B** | **B** |
|  | 50.6 | C,D | D | **A** | **B** | B,C | **B** | **A,B** |

**Supplementary Table 5**

|  |  | Pearson correlation coefficient (r) |
| --- | --- | --- |
| phospholipase MUP concentration (µM) | 10 | 0.93 |
|  | 40 | 0.77 |
|  | 60 | 0.90 |
|  | 120 | 0.86 |
| phospholipase Vmax |  | 0.66 |
| phospholipase Km |  | -0.91 |
| phospholipase efficiency |  | 0.89 |
| acyltransferase DHE concentration (µM) | 6.4 | 0.82 |
|  | 9.6 | 0.85 |
|  | 14.4 | 0.85 |
|  | 21.6 | 0.88 |
|  | 32.5 | 0.90 |
|  | 50.6 | 0.89 |
| acyltransferase Vmax |  | 0.76 |
| acyltransferase Km |  | -0.62 |
| acyltransferase efficiency |  | 0.86 |

**Supplementary Table 6**

| secondary metabolite keyword | filter 1: LCAT OR “lecithin cholesterol acyltransferase” AND (secondary metabolite keyword) | filter 2: filter 1 AND  (plant OR natural OR nature OR herbal OR nutraceutical OR bioceutical OR citrus OR vegetable OR fruit OR seed OR leaf OR stem OR supplement OR bark OR indigenous) | filter 3: manual filter |
| --- | --- | --- | --- |
| alkaloid | 3 | - | 3 |
| NPAA OR ”non protein amino acid” | 0 | - | - |
| amine | 8 | - | 0 |
| ”cyanogenic glycoside” | 0 | - | - |
| glucosinolate | 0 | - | - |
| alkamide | 0 | - | - |
| lectin | 11 | - | 1 |
| peptide or polypeptide | 174 | 17 | 4 |
| terpene | 3 | - | 3 |
| steroid | 71 | 9 | 3 |
| saponin | 4 | - | 3 |
| flavonoid | 16 | - | 16 |
| tannin | 4 | - | 4 |
| phenylpropanoid | 0 | - | - |
| lignin | 0 | - | - |
| coumarin | 4 | - | 4 |
| lignan | 1 | - | 1 |
| polyacetylene | 0 | - | - |
| ”fatty acid” | 457 | 64 | 28 |
| wax | 1 | - | 1 |
| polyketide | 0 | - | - |
| carbohydrate | 72 | 12 | 5 |
| organic acid | 2 | - | 0 |

**Supplementary Table 7**

| Secondary metabolite | DOI or reference |
| --- | --- |
| alkaloid | 10.1016/j.cbi.2014.07.008 |
|  | 10.1021/bi00227a010 |
| lectin | 10.1155/2020/2907610 |
| peptide or polypeptide | 10.3390/nu13051410 |
|  | 10.3390/nu12030610 |
|  | 10.1108/NFS-04-2016-0046 |
|  | 10.1073/pnas.77.6.3154 |
| terpene | 10.3109/13880209.2016.1168852 |
|  | 10.1016/S1995-7645(11)60167-3 |
|  | 10.1007/BF02534791 |
| steroid | 10.1248/bpb.24.713 |
|  | 10.1093/jn/120.12.1624 |
| saponin | 10.1039/c7fo01109a |
|  | Anales de la Real Academia de Farmacia 65, 327-349 (1999) |
| flavonoid | 10.1016/j.biopha.2020.110298 |
|  | 10.1016/j.biopha.2018.02.116 |
|  | 10.1016/j.jnutbio.2017.04.011 |
|  | 10.4162/nrp.2016.10.5.501 |
|  | 10.1515/jbcpp-2015-0017 |
|  | 10.1016/j.ejphar.2014.09.014 |
|  | International Journal of Pharmacognosy and Phytochemical Research 6, 758-765 (2014) |
|  | 10.1016/j.jff.2012.12.004 |
|  | 10.1080/19390210902861825 |
|  | Zhongguo Zhongyao Zazhi 33, 1064-1066, 2008 |
|  | 10.1159/000093558 |
|  | Indian Journal of Experimental Biology 41, 296-303 (2003) |
|  | 10.1016/S0378-8741(01)00361-0 |
|  | 10.1016/S0308-8146(00)00225-9 |
|  | 10.1023/A:1007965927434 |
|  | 10.3164/jcbn.19.175 |
| tannin | 10.1016/j.jff.2017.09.023 |
|  | 10.1016/j.foodres.2012.05.024 |
| coumarin | 10.1016/j.biopha.2016.10.103 |
|  | 10.1016/j.bionut.2014.02.003 |
|  | 10.2165/00003495-198733060-00002 |
|  | Artery 5, 110-116 (1979) |
| lignan | 10.31838/ijpr/2020.SP1.002 |
| wax | 10.3390/nu12010262 |
| carbohydrate | International Journal of Pharma and Bio Sciences 6, B1281-B1288 (2015) |
|  | 10.3109/19390211.2014.887599 |
|  | 10.1080/07315724.1999.10718825 |
|  | 10.1016/0885-4505(86)90106-4 |
|  | 10.1016/0021-9150(83)90075-8 |
| fatty acid | 10.1111/tpj.15050 |
|  | 10.1016/j.phymed.2020.153274 |
|  | 10.1021/acs.jafc.8b04080 |
|  | 10.3390/nu9060599 |
|  | 10.3390/nu9010082 |
|  | 10.1016/j.phymed.2013.08.027 |
|  | International Journal of Pharmacy and Pharmaceutical Sciences 6, 459-464 (2014) |
|  | 10.1155/2014/982498 |
|  | 10.4162/nrp.2013.7.4.287 |
|  | 10.1371/journal.pone.0061109 |
|  | Biochemical and Cellular Archives 13, 137-143 (2013) |
|  | 10.1111/j.1745-4514.2011.00561.x |
|  | Asian Journal of Pharmaceutical and Clinical Research 4, 154-157 (2011) |
|  | Indian Journal of Biochemistry and Biophysics 47, 104-109 (2010) |
|  | 10.1016/j.atherosclerosis.2009.03.039 |
|  | Indian Journal of Experimental Biology 47, 169-175 (2009) |
|  | General Physiology and Biophysics 27, 3-11 (2008) |
|  | 10.3136/nskkk.53.380 |
|  | 10.1159/000088935 |
|  | 10.1007/s11010-005-3194-x |
|  | 10.1016/S0024-3205(00)00665-2 |
|  | 10.1016/S0308-8146(99)00135-1 |
|  | 10.1016/S0271-5317(05)80775-4 |

**Supplementary Table 8**

| Source of compounds | DOI or reference |
| --- | --- |
| Persimmon tannin | 10.1155/2016/3424025 |
| Licorice extract | 10.1016/j.indcrop.2017.11.050 |
| Anthocephalus indicus | 10.4103/0973-7847.162110 |
| Scutellaria Baicalensis | 10.1002/jssc.201700473 |
| Emblica officinalis | 10.1002/ptr.2775 |
| Mangifera indica | Chinese Traditional and Herbal Drugs 42, 428-431 (2011) |
| Costus igneus | 10.1016/j.imu.2018.10.004 |
| Cassia auriculata | 10.1016/S0367-326X(99)00108-2 |
| Nelumbo nucifera | 10.1016/j.bmcl.2013.04.013 |
| Betula alnoides | 10.1080/0972060X.2015.1036569 |
| Garcinia cambogia | 10.1016/j.fitote.2015.02.012 |
| Solanum melongena | 10.20546/ijcrbp.2020.706.003 |

**Supplementary Fig. 1**

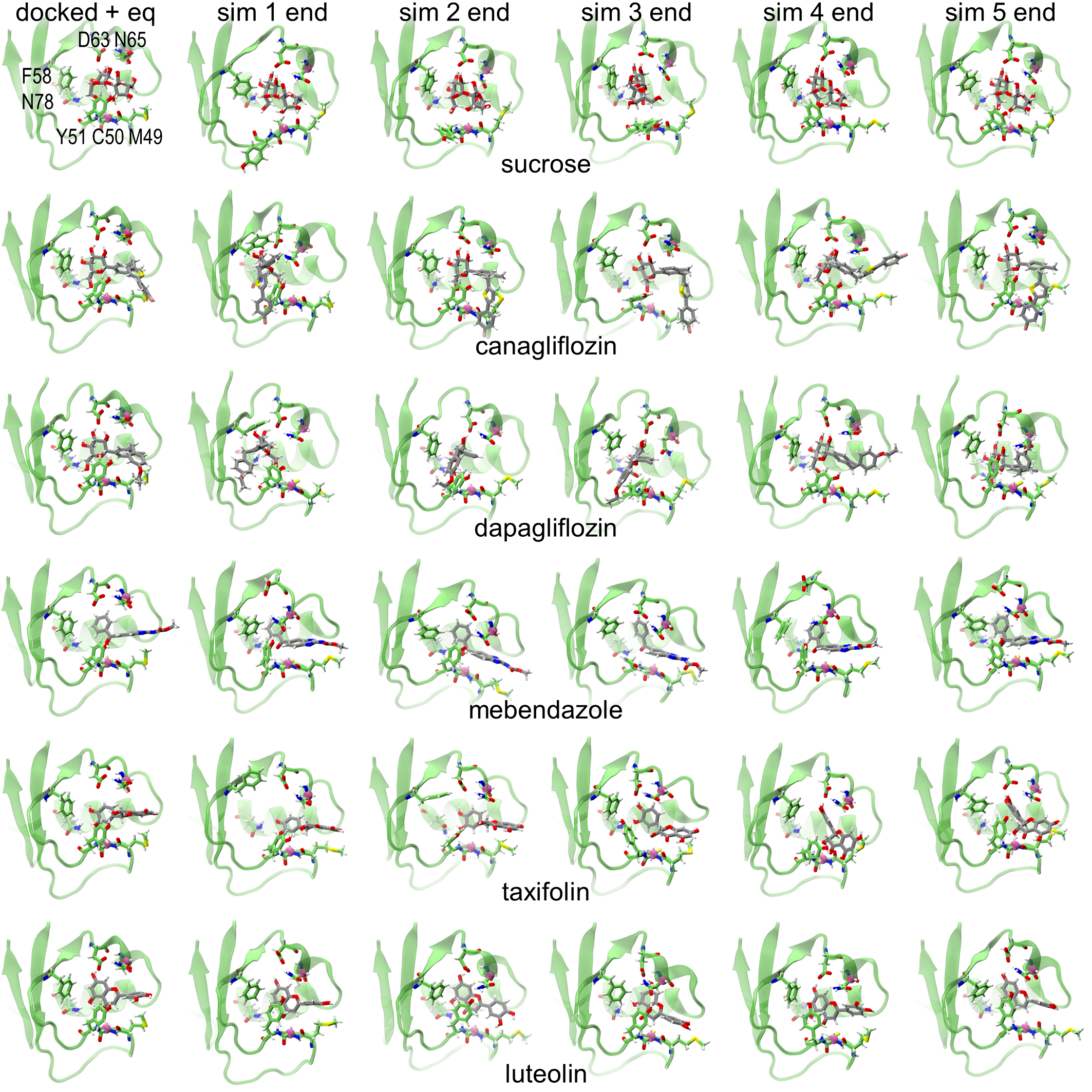

**Supplementary Fig. 2**

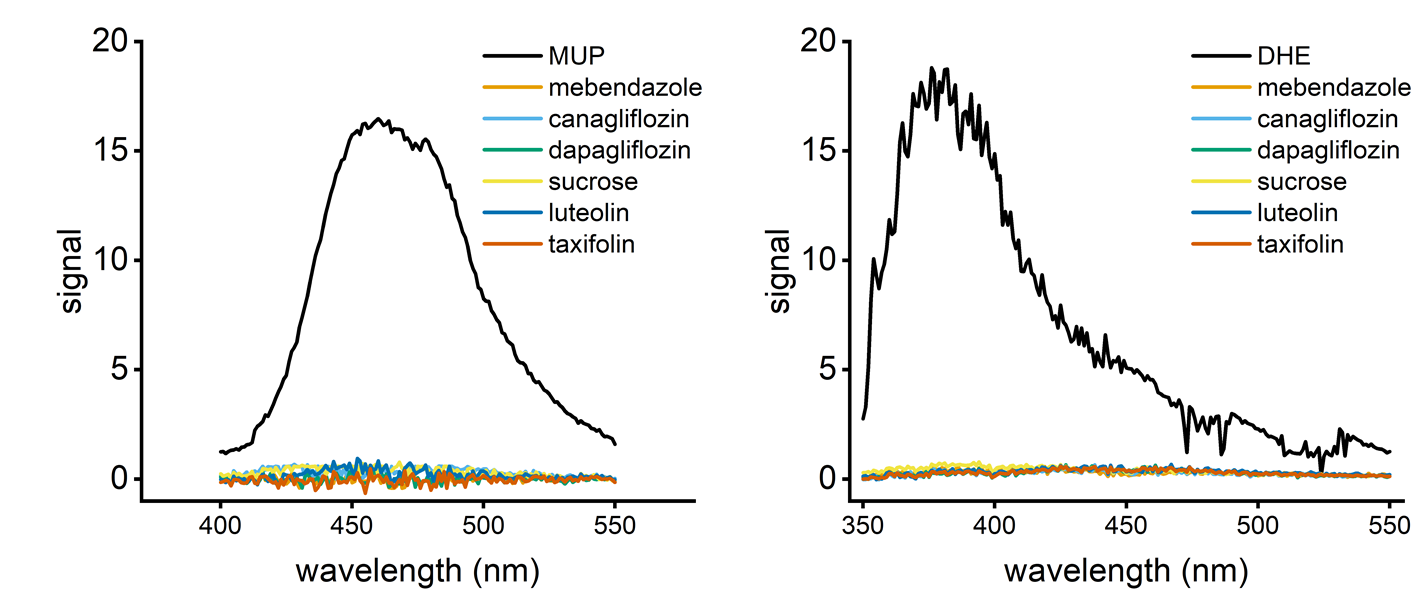

**Supplementary Fig. 3**

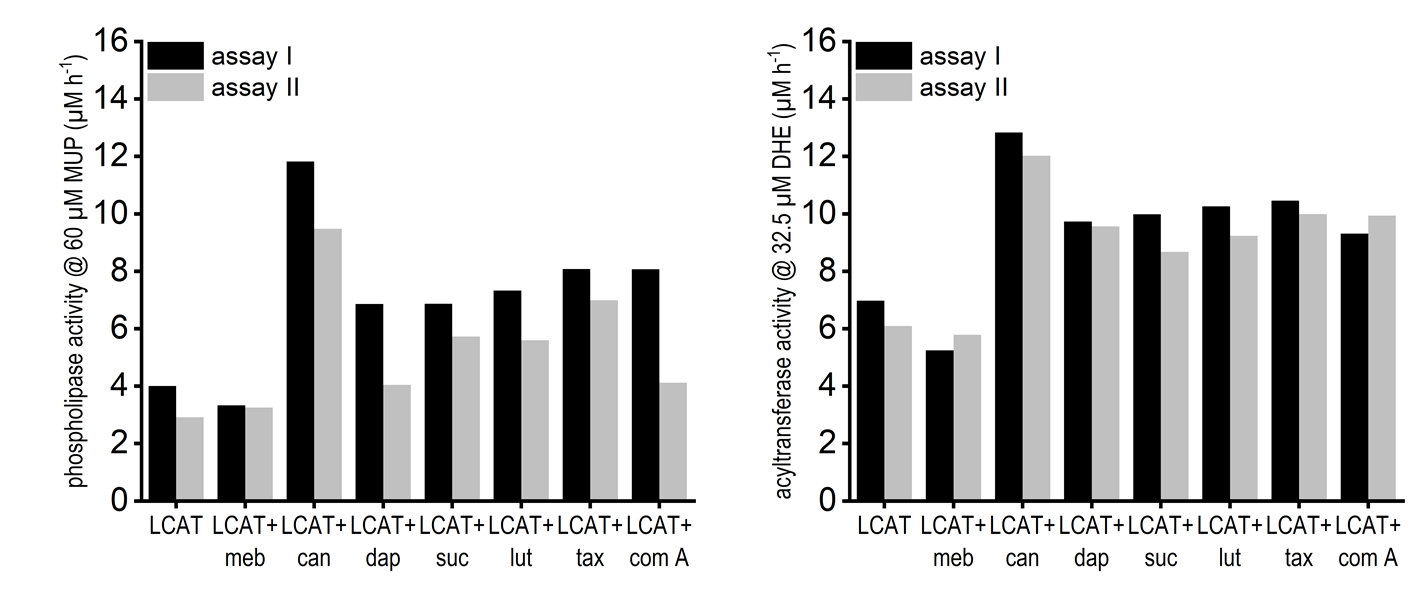
